## Supplemental Table 1 for "PET/CT targeted tissue sampling reveals virus specific dIgA can alter the distribution and localization of HIV after rectal exposure"

Supplemental Table S1: Number of tissue blocks and images per rhesus macaque in Protocol 3

| animal | tissue type | tissue blocks (total) | tissue blocks (PET selected) | sections cut/block* (total sections) | sections used/block (total sections used) | Z-stacked images/section (total images analyzed) *** |
|---|---|---|---|---|---|---|
| each | rectum | 100-130 | 4-5 | 2-3 (8-15) | 1 (4-5) | 20-22 (88-110) |
| | descending colon | | 4-5 | 2-3 (8-15) | 1 (4-5) | 20-22 (88-110) |
| | transverse colon | | 4-5 | 2-3 (8-15) | 1 (4-5) | 20-22 (88-110) |
| | mesenteric lymph nodes | | 4-5 | 2-3 (8-15) | 2 (8-10)** | 20-22 (88-110) |
| **TOTALS/animal** | | | 16-20 | 32-60 | 16-20 | **352-440** |

e; a second 2-3 section slide was created as a reserve/back up

1 $cm^2$ sections of colorectal tissue, often 2 sections per block were needed to get 22 images

*** Using a 100x lens, the 22 images (per section) were taken by simply following the linear path of the lumenal surface of the colorectal mucosa (distance between images was dictated by the bleaching caused by photoactivation). Images were shot and then only later examined and analyzed.
